# A comparison of claw removal methods on the post-release survival and claw regeneration of stone crab (*Menippe mercenaria*)

**DOI:** 10.1101/2022.10.24.513277

**Authors:** Alexandria M. Walus, Eric V. C. Schneider, Erin Parker, Candice Brittain, Iain J. McGaw, Daniel Hayes, Amber Peters, Travis E. Van Leeuwen

**Affiliations:** Department of Fisheries and Wildlife, 480 Wilson Road, Room 13, East Lansing, Michigan, USA, 48824; Cape Eleuthera Institute, PO BOX EL-26029, Rock Sound, Eleuthera, The Bahamas; Memorial University of Newfoundland, Department of Ocean Sciences, 0 Marine Lab Rd., St. John’s, Newfoundland,Canada, A1C 5S7; Fisheries and Oceans Canada, 80 East White Hills Rd., St. John’s, Newfoundland, Canada, A1C 5X1

**Keywords:** Claw removal, stone crab, autotomy, claw regeneration, fisheries

## Abstract

Commercial and recreational stone crab (*Menippe mercenaria*) fisheries primarily occur along the Gulf of Mexico and Atlantic coasts of the southeastern United States and the northeastern Caribbean. The fishery is unique in that only the crabs’ claws are retained and the animal is returned to the water alive. While the fishery is often regarded as sustainable because it is believed to exploit the crabs’ natural ability to voluntarily drop (autotomize) and regenerate lost claws, the post-release survival of de-clawed stone crabs is often low, especially when both claws are harvested. In this study, a controlled laboratory experiment was used to compare a new method of claw removal to the typical method currently used in the fishery. For the two different claw removal methods, we compared crab survival and start time to claw regeneration as a function of harvester and whether one claw or both claws were removed. Overall, we found a significant effect of the removal method, harvester, and whether one claw or both claws were removed on crab survival, but these factors did not influence the time to start of claw regeneration. Although our new method was several seconds slower in processing time than the typical method, it resulted in a 28% increase in survival (up to 92% survival throughout the study) compared to the typical method of claw removal (64% survival throughout the study). Overall, these results suggest that our new method of claw removal significantly increases post-release survival of stone crabs, and most notably does so independent of harvester and whether one claw or both claws are removed.

## 1. Introduction

Commercial and recreational stone crab (*Menippe mercenaria*) fisheries primarily occur along the Gulf of Mexico and Atlantic coasts of the southeastern United States, but smaller commercial fisheries are emerging in Belize (Bert and Hochberg, 1992) and The Bahamas (Moultrie *et al*., 2016). In The Bahamas, the stone crab fishery is currently the third largest export valued at approximately 2.4 million US dollars. These emerging fisheries primarily supplement the demand for stone crab in the United States of America (USA). Typically, crabs are captured in baited traps with fisheries regulations mandating a minimum size for claw harvest. Minimum claw-size, whether one claw or both claws can be removed, and seasonal closures vary by region and country, but best practice recommends that only a single (larger) claw be removed (Bert et al. 2016). Nevertheless, both claws are legally harvested in many commercial and recreational fisheries (Gandy *et al*., 2016, Sullivan, 1979).

While the stone crab fishery is widely recognizable as a claw-only fishery, others operate in this manner as well. In the United Kingdom, the edible or brown crab (*Cancer pagurus*) fishery removes both claws before returning the crab to the sea (Patterson *et al*., 2009). In southern Portugal, the major claw is removed from the fiddler crab (*Uca tangeri*) and is considered a local delicacy (Patterson *et al*., 2009). In the Northwest Atlantic an emerging claw-only fishery for Jonah crab (*Cancer borealis*) is being studied (Goldstein and Carloni, 2021). Claw-only fisheries are often regarded as sustainable because they are assumed to exploit the natural ability of crabs to voluntarily drop (autotomize) and regenerate their claws when damaged or threatened. Because the animal is returned alive, it is assumed to re-enter the fishery once the claws regrow and in the meantime has the ability to reproduce and provide a legacy of recruits to the fishery. However, post-release survival of de-clawed stone crabs is variable and often low, ranging between 2% and 81% in controlled laboratory conditions (Davis *et al*., 1978, Duermit *et al*., 2015, Simonson and Hochberg, 1986) and between 37% and 59% in the wild (Gandy *et al*., 2016). The large variation in survival is due to a myriad of factors that influence survival after claw removal and release. These include, but are not limited to, water temperature (Gandy *et al*., 2016), break location (Gandy *et al*., 2016), wound size (Duermit *et al*., 2015, 2017), and whether both claws are removed (Davis *et al*., 1978, Duermit *et al*., 2015, Gandy *et al*., 2016; Orrell *et al*., 2019).

Typically, claw removal is accomplished by exerting enough downward force to the fully extended cheliped to break the claw cleanly along the autotomy plane at the basi-ischium, located between the coxa at the base of the leg and merus. If done correctly, it allows for the formation of a hypodermal diaphragm (Savage and Sullivan, 1978) which minimizes hemolymph loss at the release site (Savage and Sullivan, 1978) and results in high survival of the crab following release (Duermit *et al*., 2015, 2017). However, this is often not the case, even for experienced commercial harvesters. In the 2011 Florida commercial stone crab fishery, an average of 31% of the claws observed by Florida Fish and Wildlife Conservation Commission samplers in fish houses statewide showed evidence of poor breaks (FWC, 2011). Therefore, given that poor breaks often result in a fatal wound for the crab and 1329 metric tons (2 929 943 lbs) of claws were harvested in the 2011 commercial fishery, it is not inconceivable that approximately 11.7 to 17.6 million crabs (assuming 4 to 6 claws per pound and one claw removal per crab) had their claws harvested in the fishery and approximately 3.6 to 5 million crabs may have died as a result of claw removal (number of crabs * 0.31 i.e. proportion of crabs in the fishery observed to have poor breaks). Further, while the number of participants or the magnitude of the harvest is unknown for the recreational fishery, participants can fish five traps per person or catch the crabs while diving. Therefore, given that novices removing claws in the recreational fishery will likely have higher mortality than experienced commercial harvesters, the number of mortalities in the recreational fishery is also likely high. Although the number of crabs harvested in the recreational fishery is likely magnitudes lower than in the commercial fishery, this may still account for unnecessarily high mortality rates that affect crab stocks.

Lastly, despite the importance of increasing survival in any fishery, decapod crustacean welfare awareness in the past decade has been increasing, with multiple review articles covering the topic (e.g. Stoner et al 2012; Sneddon et al 2014, Diggles 2019). In 2021, the Government of the United Kingdom backed a bill that recognizes decapod crustaceans as sentient beings (Animal Welfare Bill, 2021: Birch et al, 2021). This policy is now spreading across Europe and is expected to be implemented by other Nations. Thus, it is expected that this will have far-reaching impacts not only on the use of decapods in scientific research, but their general welfare in commercial and recreational fisheries.

In this study, a controlled laboratory experiment was used to compare a new method of claw removal to the typical method used in this fishery. For the two different claw removal methods, we compared the survival and start time for claw regeneration as a function of harvester and whether one claw or both claws were removed.

## 2. Materials & methods

### 2.1 Animal collection and husbandry

One hundred and eighteen stone crabs (mean mass ± SD = 276.2 ± 65.6 g; mean carapace width ± SD = 95.1 ± 9.3 mm; right propodus length mean ± SD = 74 ± 9.5 mm; left propodus length mean ± SD = 69.57 ± 8.7 mm) with both claws intact were sourced from a stone crab harvester in Hatchet Bay, Eleuthera, The Bahamas. The crabs were covered in seawater-soaked towels and transported in large coolers to the wet lab facility at the Cape Eleuthera Institute, Eleuthera, The Bahamas. The crabs were acclimated for 21 days prior to experimentation in two large circular aerated tanks (3.7m diameter, 1m water height) supplied with flow through seawater at a temperature of 26.3 (± 1.2 SD) °C and a salinity of 36 (± 1 SD) psu. The crabs were fed pieces of conch and fish offal *ad libitum* every other day. PVC pipes were added to the tanks to provide enrichment and shelter and to help minimize agonistic behaviors among crabs. Holding tanks were cleaned bi-weekly.

### 2.2 Claw removal and experimental treatments

Five days prior to claw removal trials, all crabs (female = 58, male = 60) were tagged using a small piece of numbered waterproof paper (Rite-In-The-Rain; JL Darling LLC, Washington, USA) that was glued (Krazy Glue; Ohio, USA) to the carapace, allowing identification of individual crabs throughout the experiment. To determine the effect of claw removal method, harvester, and whether one claw or both claws were removed on the post-release survival and start of claw regeneration time, crabs were randomly assigned to one of nine treatment groups (Table 1). Briefly, these included two people, a commercial fisher (experienced harvester) and a novice fisher (novice harvester) removing one or two claws using either the typical method used in the fishery or the new method. The commercial harvester was a stone crab fisher on the island of Eleuthera with > 10 years’ experience, whereas the novice harvester was a researcher with limited claw removal experience but was familiar with handling stone crabs. The typical method was conducted by applying a downward force to the crabs fully extended cheliped until the claw broke along the autotomy plane at the basi-ischium, located between the coxa at the base of the leg and merus (Figure 1). The new method was conducted by puncturing the arthrodial membrane (the soft joint, or unsclerotized cuticle) between the carpus and the merus with a marlin spike (5 cm long X 2mm diameter metal spike often used to splice rope and repair sails). This was accomplished by pushing the crab towards the mounted spike that was horizontally clamped to a table, and slowly rotating the crab back and forth until the claw released. This resulted in the induced release of the claw (autotomy) distal to the coxa (Figure 2). Following claw removal, all crabs were haphazardly released into one of three flow-through circular tanks (3.7m diameter, 1m water height) and held under the same conditions and feeding regime described above for acclimation. Data recorded included: left and right claw length, left and right claw weight (for the claw removal group and depending on which claw was removed), carapace width, incorrect break occurrence (claw removals that resulted in visible signs of damage to the coxa), sex, and the total amount of time it took harvesters to remove claws per claw-removal treatments.

**Table 1.** Breakdown of treatments, treatment abbreviations and sample size of crabs used in the experiment to compare a new method of claw removal to the typical method of claw removal. For the two different claw removal methods, we compared the survival and start time for claw regeneration as a function of harvester (experienced and novice) and whether one claw or both claws were removed.

| Treatment | Abbreviation | Sample size (n) |
| --- | --- | --- |
| Experienced harvester-1 claw removal-Typical method | E-1-T | 14 |
| Experienced harvester-2 claw removal-Typical method | E-2-T | 13 |
| Experienced harvester-1 claw removal-New method | E-1-N | 14 |
| Experienced harvester-2 claw removal-New method | E-2-N | 14 |
| Novice harvester-1 claw removal-Typical method | N-1-T | 14 |
| Novice harvester-2 claw removal-Typical method | N-2-T | 14 |
| Novice harvester-1 claw removal-New method | N-1-N | 7 |
| Novice harvester-2 claw removal-New method | N-2-N | 14 |
| Handled but no claw removal | Control | 14 |

**Figure 1.**
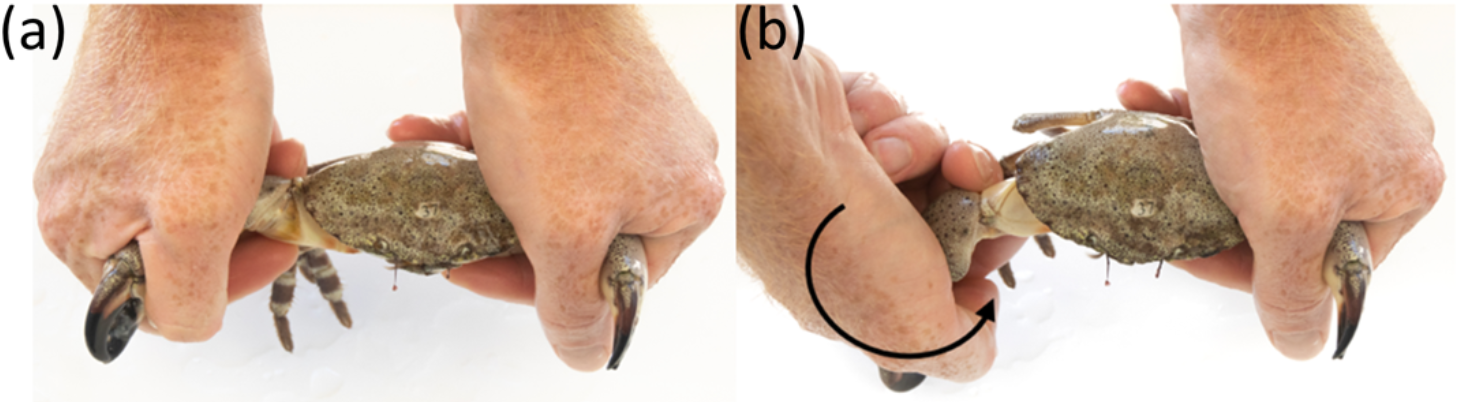
A photo series of the typical recommended method of claw removal for stone crabs (*Menippe* spp.) and other claw-only fisheries. This method uses (a) two hands to hold the crabs claws, and (b) a downward force applied to the crabs fully extended cheliped to break the claw along the autotomy plane at the basi-ischium, located between the coxa at the base of the leg and merus.

**Figure 2.**
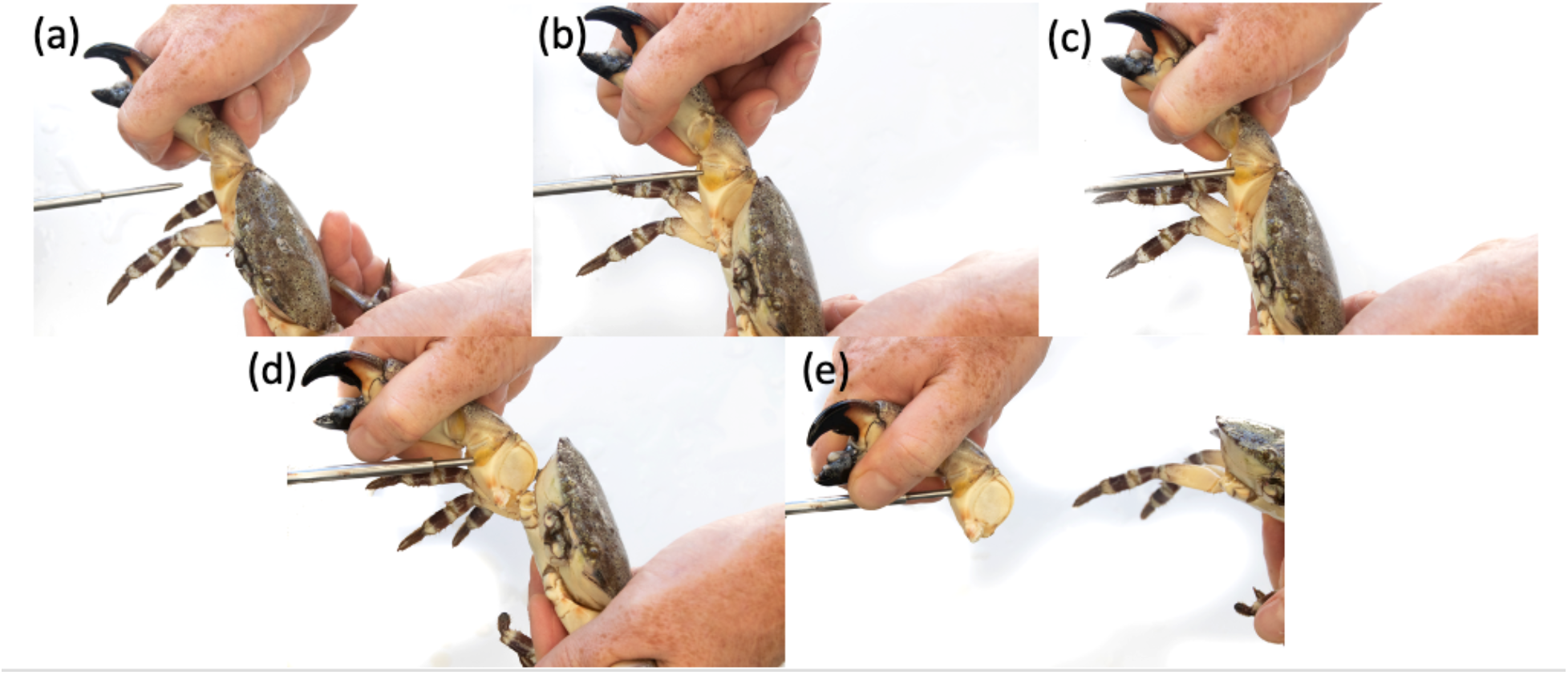
A photo series of our new method of claw removal for stone crabs (*Menippe* spp.) and potentially other claw-only fisheries. This method uses a marlin spike (a) to puncture the articular membrane (the soft joint, or unsclerotized cuticle between hard sclerotized parts of the exoskeleton) between (b, c) the carpus and the merus and (d) rotating the implement towards the crab’s body and inducing (e) the voluntary drop (autotomy) of the claw distal to the coxa.

### 2.3 Survival and time to start of claw regeneration

Following claw removal, crabs were checked daily for 35 days and then weekly for the next 25 days to assess mortality and signs of claw regeneration. If mortality occurred, the individual tag identification number and the number of days to the mortality event from the time of claw removal was recorded and the individual was removed from the experiment. The start of claw re-growth was assessed as the presence or absence of a small protrusion (precursor to the limb bud) forming at the removal site of the coxa where the claw (cheliped) was removed. If a crab was observed to have started claw regeneration, the individual tag identification number, and the number of days to the start of claw regeneration from the time of claw removal was recorded.

Following experiments all remaining crabs were released back into the wild. All work was carried out under The Bahamas Department of Marine Resources permit number MAMR/FIS/2/12A/17/17B.

### 2.5 Calculations and statistical analyses

We used a Cox proportional hazards (CPH) analysis using the R packages “survival” (Therneau, 2015) and “survminer” (Kassambara, 2021) to examine potential effects of removal method (typical or new), harvester, whether one claw or both claws were removed, sex and carapace width on the post-release survival of stone crabs. Additionally, a Kaplan-Meier (KM) analysis using the same R packages as above, and days to the event (dead or alive) was used to determine survival rate estimates over the course of the study (1 to 35 days) across removal methods, harvester, and claw removal treatments. To compare the cumulative effects of removal methods (i.e. differences among individual treatments; Table 1), harvester and claw removal treatments on survival (i.e. 35 days following claw removal), we used a Kruskal Wallis test to compare survival among individual treatments. To determine the effect of incorrect claw breaks on survival and to compare the number of incorrect breaks that occurred among treatments, we used a Spearman’s-Rank correlation test and a Kruskal Wallis test, respectively. Procedure times of the various treatments was calculated by dividing the number of crabs that were de-clawed in a treatment by the total claw removal time for that treatment. While procedure times for individual crabs per treatment would have been ideal, the addition of this data was opportunistic but nevertheless important enough to be included for context in highlighting potential differences in procedure times between the typical removal method used in the fishery and the new method. This is an important consideration in the feasibility and adoption with any proposed modification to a fishery procedure, where excess time often results in greater costs and thus low support for the adoption of the proposed change. Lastly, to examine the effect of claw removal method, harvester, and whether one claw or both claws were removed on the time to the start of claw regeneration we again used a Kruskal Wallis test.

## 3. Results

### 3.1 Survival, break quality, and procedure times

There was a significant effect of claw removal method (CPH = −1.90, z = −3.40, p < 0.001), whether one claw or both claws were removed (CPH = 0.96, z = 2.14, p = 0.03), and between harvesters (CPH = 0.90, z = 2.16, p = 0.03), on the post-release survival of crabs. However, there was no significant effect of sex (CPH = 0.31, z = 0.63, p = 0.53) or carapace width (CPH = 0.003, z = 0.14, p = 0.89). Overall, survival for the control treatment (handled but no claw removal) was 93% and crabs from the one claw removal treatments had higher survival (84%) than crabs from the two claw removal treatments (70%). Survival of all crabs with claws removed via both methods by the more experienced harvester was 84% whereas survival of all crabs with claws removed via both methods by the novice harvester was 67%. Survival using our new method was 92% whereas those removed across the typical method treatments was 63% (Figure 3A). Therefore, our new method resulted in a 29% increase in survival relative to the typical method and only a 1% decrease in survival relative to the control. Finally, there was a significant cumulative effect of removal method (i.e. treatment effect), harvester and whether one claw or both claws were removed on the survival of crabs (Kruskal Wallis = 31.45, df = 8, p < 0.001; Figure 3B). Crabs with both claws removed by the novice harvester using the typical method (N-2-T) had the lowest survival at 43% (6 / 14), while crabs with one claw removed by the experienced harvester using our new method (E-1-N) had the highest survival at 100% (14 / 14; Figure 3B).

**Figure 3.**
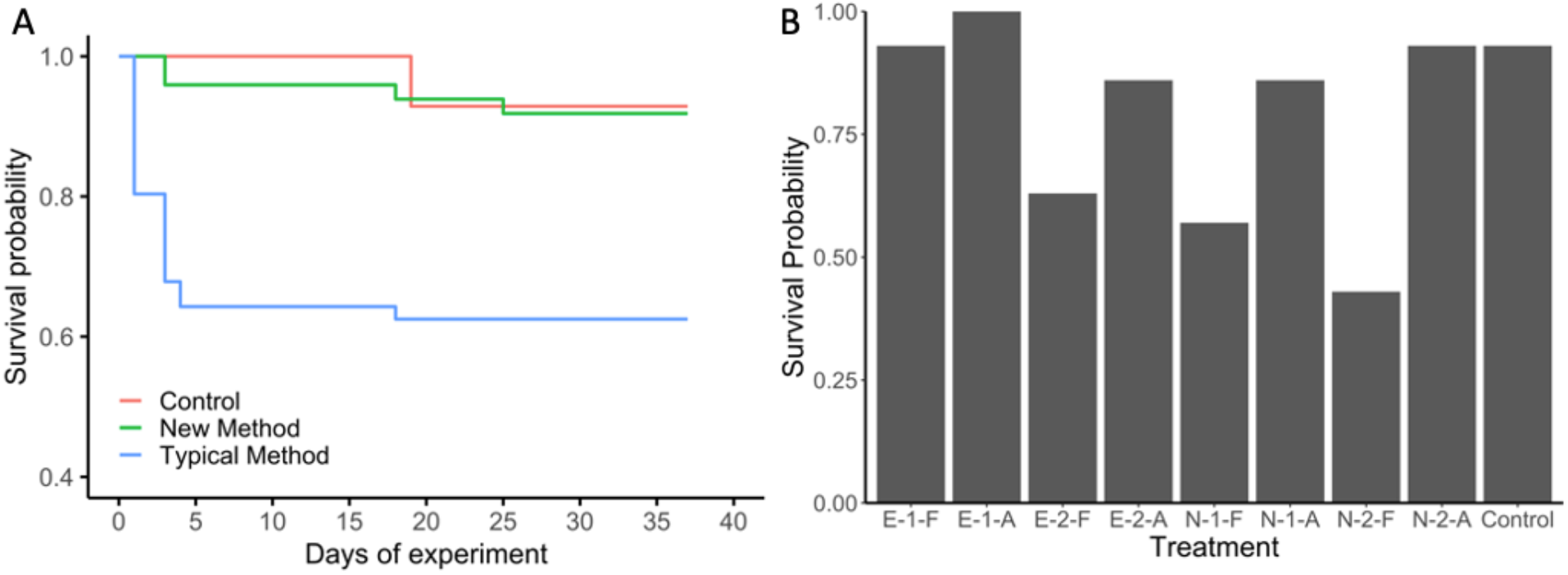
(A) Kaplan-Meier survival curves for crabs used in the experiment to compare a new method of claw removal to the typical method of claw removal used in the fishery. (B) Mean survival probability of crabs across individual treatments [(Expert (E) or Novice (N) – 1 claw removed (1) or 2 claws removed (2) – typical method (T) or new method (N)]. Mean water temperature and salinity for the study ± SD was 26.3 ± 1.2°C and 36 ± 1 psu.

As expected, the number of incorrect claw breaks that occurred among treatments was negatively correlated with post-release survival (ρ_7_ = −0.83, p = 0.005; Figure 4). Crabs with both claws removed by the novice harvester via the typical method (N-2-T) resulted in the greatest number of incorrect claw breaks with 64% (9 / 14; Kruskal Wallis = 32.42, df = 7, p < 0.001; Figure 4) of claw removals resulting in incorrect breaks and ultimately the lowest survival of all treatments at 43% (6 / 14). However, the E-1-T, E-1-N and N-1-N treatments all resulted in clean claw breaks during all removals.

**Figure 4.**
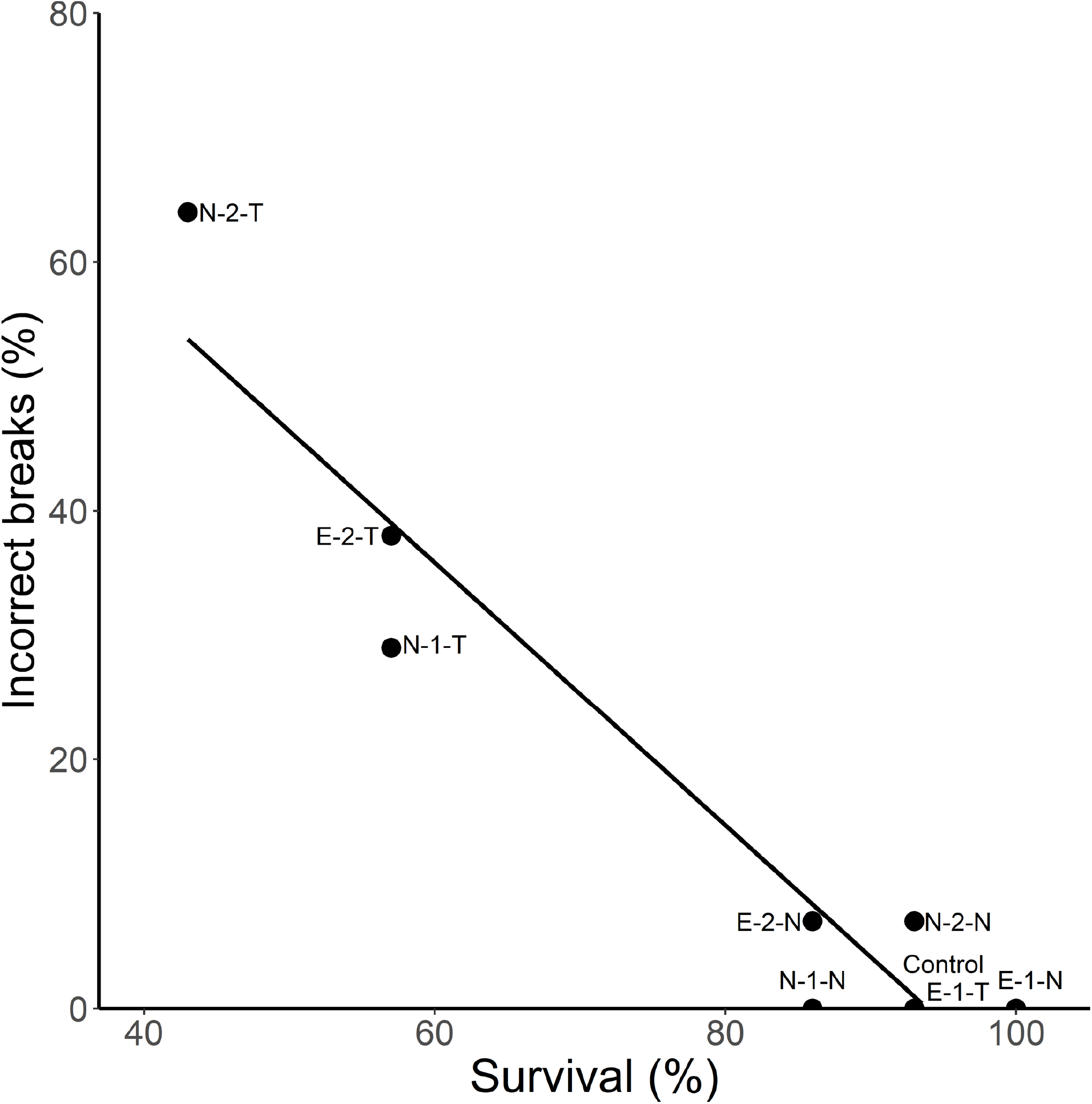
The relationship between incorrect claw breaks (claw removals that resulted in a wound) and survival for the various treatments of the study [(Expert harvester (E) or Novice harvester (N) – 1 claw removed (1) or 2 claws removed (2) – Typical method (T) or New method (N)].

The removal time for the one claw removal treatments and the experienced harvester treatments were the fastest and took an average of 4.6 and 8.4 seconds per crab, respectively. The two claw removal treatments and the novice harvester treatments were the slowest and on average took 10.7 and 8.4 seconds per crab, respectively. The new method treatments on an average were slower and took 11.4 seconds per crab to complete, whereas the typical method treatments took an average of 4.7 seconds to complete. Overall, the experienced harvester removing one claw using the typical method treatment (E-1-T) was the fastest, with an average of 1.1 seconds per crab while the experienced harvester removing two claws using our new (E-2-N) method was the slowest, with an average of 16.4 seconds per crab.

### 3.2 Start of claw regeneration time

There was no significant effect of removal method (Kruskal-Wallis = 0.92, p = 0.33), harvester (Kruskal-Wallis = 0.0005, p = 0.98), and whether one claw or both claws were removed (Kruskal-Wallis = 0.12, p = 0.73), on the start of claw regeneration time. Among treatments, crabs took approximately 40 days for the start of claw regrowth to be visually observed.

## 4. Discussion

The results of our study showed that not only does our new method increase survival, but it does so irrespective of harvester and whether one or both claws are removed. Survival among our new method treatments was 92% (99% relative to the control) and is among the highest reported in the literature. For example, Gandy *et al*. (2016), in a field-based experiment, found that overall survival of crabs following claw removal was 48% (59% for one claw removals and 37% for two claw removals). In the laboratory, Davis *et al*. (1978), found an overall survival of 63% (72% survival for one claw removals and 53% survival for two claw removals), whereas Simonson and Hochberg (1986), found an overall survival of 80% for a single claw removal. While some differences exist in environmental and experimental conditions among studies, the key difference between survival reported in the literature and our study is the method used to remove the claw. Our new method produced a clean break across the autotomy plane 96% of the time. In contrast, the typical method only produced a clean break 67% of the time with 33% of cases resulting in damage or hemolymph loss at the release site (Savage and Sullivan, 1978) and subsequent death of the crab one to five days post-release.

While the typical method of claw-removal in the stone crab fishery is believed to exploit the natural ability of crabs to voluntarily drop (autotomize) their claw, it is clear that this is not always the case, and damage can regularly occur across the autotomy plane. This damage may occur even after accounting for harvester experience and whether both claws are removed and is the most likely contributor to the poor survival in the fishery. Common injuries include breaking the claw into the merus side of the basi-ischium segment, or breaking the claw and exposing the body cavity (Simonson and Hochberg, 1986, Gandy *et al*., 2016). Both of these injuries have the potential for loss of hemolymph and also leave an open surface for bacterial infection to occur (Simonson and Hochberg, 1986, Gandy *et al*., 2016). The higher incidence of damage that occurred during removal of the second claw likely occurs because stone crabs are reliant on their claws for protection and feeding: therefore, once a claw has already been removed the resulting level of perceived impairment by the crab increases and it requires more force to remove subsequent limbs (Robinson *et al*., 1970), which further increases likelihood of a damaging claw break using the typical method.

Interestingly, the effect of claw removals across treatments was for the most part only observed when comparing among the typical method treatments (i.e. experienced harvester-1 claw removal-typical method (E-1-T) and experienced harvester-2 claw removal-typical method treatments (E-2-T) to the novice harvester-1 claw removal-typical method (N-1-T) and novice harvester-2 claw removal-typical method treatments (N-2-T)) and not among our new method treatments (i.e. experienced harvester-1 claw removal-new method (E-1-N) and experienced harvester-2 claw removal-new method treatments (E-2-N) to the novice harvester-1 claw removal-new method (E-1-N) and novice harvester-2 claw removal-new method treatments (N-2-N)). This result further suggests that our new method is likely what resulted in the higher survival as result of the much lower incidence of damage across the break plane and much higher success of eliciting the behavioral autotomy response of the crab that occurs when the claw is damaged or the animal is threatened. In fact, the novice harvester-2 claw removal-new method treatment (N-2-N) had a 7% higher mean survival than the experienced harvester-2 claw removal-new method treatment (E-2-N) and although the experienced harvester-1 claw removal-new method treatment (E-1-N) had a slightly higher mean survival than the experienced harvester-2 claw removal-new method treatment (E-2-N), the novice harvester-2 claw removal-new method treatment (N-2-N) had a slightly higher survival than the novice harvester-1 claw removal-new method treatment (N-1-N). While it is difficult to determine the number of claw removals required before maximum survival of the typical method is achieved by a harvester, it is clear that there is a learning curve to conducting the typical method as significant variation in survival was found between harvesters. For example, survival following a single claw removal by the experienced harvester using the typical method was 36 % higher (93 % over the course of the study) than the same treatment conducted by the novice harvester (57 % over the course of the study). However, both harvesters took similar times to remove claws using the new method. This is not unexpected, as both harvesters were new to this procedure. While the time to perform the new method appears longer than the typical method, we expect with practice, that the time would be reduced and claw harvesting efficiency improved. Even if the new method is not as rapid, crab survival rates would increase and likely improve the future catch rates and sustainability of the fishery and undoubtedly general welfare in commercial and recreational stone crab fisheries. However while the results are promising, this experiment was carried out under controlled laboratory conditions and further studies employing this method in the field in combination with perhaps mark and recapture should be carried out to validate the transferability of the laboratory study presented here to the field.

There was no difference in the start time for claw regeneration when comparing the new method of claw removal to the typical method. This is not unexpected because the crabs with claw breaks during removal subsequently died and thus the crabs that regenerated claws were predominately those that had clean claw breaks, irrespective of the method used to remove the claws. In the present study, crabs took approximately 40 days to show the first signs of claw regrowth, but it usually takes an additional 2 to 4 years for these claws to reach harvestable size, depending on regional size regulations (Savage and Sullivan, 1978). Therefore, animals that can be legally harvested are estimated to be between 3 to 4 years old (Savage and Sullivan, 1978), so for an animal that has an average lifespan of approximately 8 years, they may not be able to regenerate harvestable claws again to re-enter the fishery (Savage and Sullivan, 1978). However, population-level reproductive benefits may still be incurred by increasing survival of crabs after harvest, despite known negative effects of claw loss and regeneration on growth, spawning fitness including fecundity, and mating success (Bender, 1971, Davis *et al*., 1978, Savage and Sullivan, 1978, Juanes and Smith, 1995, Wilber, 1995, Hogan and Griffen, 2014, Duermit *et al*., 2015).

Nevertheless, the tool required to conduct the new method of claw removal is simple and easily purchased or constructed, thus it can be easily taught to recreational and commercial harvesters as a way to improve survival and the overall sustainability of the fishery.

## Acknowledgements

We would like to thank the staff, interns, and visiting researchers of The Cape Eleuthera Institute, including Nick Higgs, Liberty Boyd, Kennedy Wall and Cameron Raguse as well as John Cartwright for providing the crabs used in the study. The study was supported through direct and indirect donor support to the Cape Eleuthera Institute and in part by the Robert C. Ball and Betty A. Ball Fisheries and Wildlife Fellowship to A. Walus. This is contribution #4 of the Exuma Sound Ecosystem Research Project.

